# Arousal threshold reveals novel neural markers of sleep depth independently from the conventional sleep stages

**DOI:** 10.1101/2024.08.09.607376

**Authors:** Dante Picchioni, Fan Nils Yang, Jacco A. de Zwart, Yicun Wang, Hendrik Mandelkow, Pinar S. Özbay, Gang Chen, Paul A. Taylor, Niki Lam, Miranda G. Chappel-Farley, Catie Chang, Jiaen Liu, Peter van Gelderen, Jeff H. Duyn

## Abstract

Reports of sleep-specific brain activity patterns have been constrained by assessing brain function as it related to the conventional polysomnographic sleep stages. This limits the variety of sleep states and underlying activity patterns that one can discover. The current study used all-night functional MRI sleep data and defined sleep behaviorally with auditory arousal threshold (AAT) to characterize sleep depth better by searching for novel neural markers of sleep depth that are neuroanatomically localized and temporally unrelated to the conventional stages.

Functional correlation values calculated in a four-min time window immediately before the determination of AAT were entered into a linear mixed effects model, allowing multiple arousals across the night per subject into the analysis, and compared to models with sleep stage to determine the unique relationships with AAT. These unique relationships were for thalamocerebellar correlations, the relationship between the right language network and the right “default-mode network dorsal medial prefrontal cortex subsystem,” and the relationship between thalamus and ventral attention network. These novel neural markers of sleep depth would have remained undiscovered if the data were merely analyzed with the conventional sleep stages.

**Significance Statement:** The original classification of sleep into distinct stages used behavioral characteristics.

With the proliferation of new techniques, the first experiments might have been to perform correlations with arousal threshold. These experiments have never been performed, either with functional MRI or with any other modern technique. The amount of communication between brain regions as measured by all-night functional magnetic resonance imaging was correlated with arousal threshold. This revealed novel neural markers of sleep depth that would have remained undiscovered if the data were merely analyzed with the conventional stages. This expands our understanding of sleep and its functions beyond the constraints imposed by the conventional stages.

## Introduction

Sleep research and sleep medicine have benefited from the advent of electroencephalography (EEG) but have also suffered from an overreliance on the conventional EEG sleep stages (e.g., Decat et al., 2022; Himanen & Hasan, 2000; Kay et al., 2017). Functional magnetic resonance imaging (MRI) can localize brain activity with much better spatial fidelity than scalp EEG. Neuroanatomical localization is important because it enables the detection of local sleep (e.g., Jang et al., 2022) and facilitates the linkage of the various brain states that occur during sleep to the functions of sleep (Maquet et al., 1997). Functional MRI can thus provide a parallel avenue of research to complement EEG sleep research. Sleep stages are defined with polysomnography (PSG), but the term EEG will be used because it is the most important measure in PSG.

The conventional sleep stages are defined with EEG, but sleep is not. Sleep is defined by behavioral characteristics (Flanigan et al., 1973; Kleitman, 1963; Piéron, 1913). When EEG was invented, researchers (Blake & Gerard, 1937) discovered a strong correlation between scalp EEG 0.5-3.0 Hz waves and auditory arousal threshold (AAT). Thus, the EEG is a *surrogate* for the behavioral definition of sleep. The continued importance of the behavioral definition is best demonstrated when investigators must define sleep in novel model organisms or at early developmental stages (e.g., Blumberg et al., 2005; Hendricks et al., 2000). When other techniques were invented to measure the brain during sleep, the first experiments should have been to perform correlations with arousal threshold. These experiments have never been performed, either with functional MRI or with any other modern technique.

Many have measured brain function during sleep with modern techniques (Maquet, 2000; Picchioni et al., 2013; Tarokh et al., in press). However with few exceptions, these studies reported sleep-specific brain activity/functional correlation patterns constrained by the conventional sleep stages. This limits the variety of sleep states and underlying activity patterns that one can discover.

If undiscovered spatiotemporal patterns of activity/functional correlations exist during sleep, one would make the following prediction: when studying one conventional stage, investigators should observe different brain and behavioral results within this stage or between new subdivisions for it. This has been found with arousal thresholds between tonic and phasic stage rapid eye movement (REM) sleep (Ermis et al., 2010; Price & Kremen, 1980), with unique theta fluctuations across stage REM sleep in mice and humans accompanied by uniquely patterned cortical waves in mice (Bueno-Junior et al., 2023; Dong et al., 2022), with unique thalamic and cortical substates in stage REM sleep that do not correspond with the conventional tonic versus phasic REM substates (Bastuji et al., 2024), with changes in autonomic nervous system tone within stage nonrapid eye movement (NREM) 2 sleep (Brandenberger et al., 2005), with a new NREM sleep stage characterized by ponto-geniculo-occipital waves occurring immediately before the transition to stage REM sleep in rats (Datta, 2000), and with differing neuroimaging brain activity/functional correlation patterns within the same stage (Guo et al., 2023; Picchioni et al., 2008; Stevner et al., 2019; Watanabe et al., 2014; Wehrle et al., 2007; Yang et al., 2024). A conventional sleep stage is defined as a pattern of scalp electrical activity associated with arousal thresholds higher than wakefulness. If this definition is expanded to include any neuroanatomically specific pattern of brain activity/functional correlations available from advanced methods, the above results are unsurprising and warrant a continued and vigorous search for new sleep states unconstrained by the conventional sleep stages.

The current study used all-night functional MRI sleep data (Moehlman et al., 2019) and defined sleep behaviorally with AAT to characterize sleep depth better by searching for novel neural markers of sleep depth that are neuroanatomically localized and temporally unrelated to the conventional stages. Functional correlation values calculated in a four-min time window immediately before the determination of AAT were entered into a linear mixed effects model, allowing multiple arousals across the night per subject into the analysis, and compared to models with sleep stage to determine the unique relationships with AAT.

## Materials and Methods

### Overview

A description of the general method, its feasibility, and its validation has been published, along with a detailed description of the subjects, screening procedures, home-monitoring period, adaptation night, functional MRI data collection, simultaneous EEG data collection, and sleep staging (Moehlman et al., 2019). Briefly, all-night functional MRI data were analyzed from 12 subjects. The basic inclusion screening criteria were: able to understand the procedures and requirements and give informed consent, fluent in English, 18-34 years of age, and in good general health. The basic exclusion criteria were neurological disorder; seizures; central nervous system surgery; current diagnosis of any psychiatric disorder; lifetime diagnosis of a psychotic, bipolar, or depressive disorder; pregnant or nursing; severe medical problem such as uncontrolled hypertension; contraindications for MRI; or subjective hearing problems. The following screening procedures were also used: standard questionnaires to detect common sleep disorder (e.g., Insomnia Severity Index; Bastien et al., 2001); a custom questionnaire designed to detect night shift work, regular use of drugs known to affect sleep, and relatively rare sleep disorders where a single item can be effective (e.g., cataplexy in Narcolepsy Type 1); a medical history evaluation and physical examination; an objective audiology examination; structural MRI brain scans; a 14-day home-monitoring period with wrist actigraphy immediately before sleeping in the scanner; and an adaptation night. No sleep deprivation was used.

### Procedure

Sleep scans began at approximately 23:00 and ended at approximately 07:00. These times corresponded to the times during which subjects were asked to sleep during the home-monitoring period to capitalize on the circadian propensity for sleep. These times were not customized to each subject’s chronotype because these times guaranteed that other users in the MRI facility would not make overlapping scan time requests and because if subjects exhibited an extreme chronotype, they would not be good candidates for the study. Data were acquired continuously throughout the night with the exceptions of equipment problems, break requests from the subjects, and experimental arousals with AAT determinations. Subjects were given the following instruction. “Please try to sleep. We will attempt to arouse you throughout the night with sounds. If you cannot sleep, that is fine, but please continue to try to sleep until you hear the sound.

When you hear the sound, please say’I’m awake.’ You must say these words exactly. You can be removed from the scanner any time that you want. If you need something, just start talking. If you do so and we do not respond, you can use the emergency squeeze bulb.“

An active noise cancellation system was used (OptoActive, OptoAcoustics) to attenuate the scanner acoustic noise and deliver the auditory tones. Sets of tones were delivered approximately eight times throughout the night to collect AAT as a behavioral measure of sleep depth. Tones were generated and delivered with Presentation (NeuroBehavioral Systems). Tones were 1.25 kHz, which was chosen to be similar to the 1.0 kHz frequency typically used while avoiding harmonics of the 0.5-kHz functional MRI acoustic noise fundamental frequency. Each tone set consisted of a sequence of five 0.1-s stimuli, separated by 0.5-s intervals.

The method of limits was used. A tone was delivered and, if the subject did not arouse, another tone was delivered 21 s later with a 5 dBA increase in volume. This continued until subjects gave a specific, arranged verbal response: “I’m awake.” If subjects did not arouse to 120 dBA, a threshold of 125 dBA was assigned as the finding, an approach that mirrors prior work.

The intensity of the first tone was customized for each subject and each night. This was determined during wakefulness before lights-off by administering a hearing test with subjects in the MRI while replicating the same experimental procedures that would occur during sleep, including the use of insert ear plugs and functional MRI data collection. AAT was defined as dBA above this waking perceptual threshold.

We used eight randomly scheduled arousals, prevalence-weighted towards stage NREM 3 sleep. This number was chosen to be in the range (2-16) of the number of arousals used in previous studies. Using randomly scheduled arousals to move beyond the conventional EEG sleep stages is novel but not unprecedented. Others have used randomly scheduled arousals, for example, in the context of studying the high-density EEG correlates of dreaming (Siclari et al., 2017) and summarized the importance of this approach by saying investigators need “to move beyond the REM-NREM sleep dichotomy and beyond traditional sleep staging” (Nir & Tononi, 2010, p. 95).

Functional MRI data were collected as previously described (Moehlman et al., 2019) with a repetition time = 3 s and a resolution = 2.5 × 2.5 × 2.0 mm^3^. Recording of peripheral signals was synchronized with functional MRI data collection with a volume trigger provided by the scanner. Concurrently acquired peripheral signals included a chest respiratory effort belt (to calculate respiratory flow rate) and a finger skin photoplethysmograph (to calculate peripheral vascular volume). They were acquired using AcqKnowledge with TSD200-MRI and TSD221-MRI transducers and an MP150 digitizer sampling at 1.0 kHz (Biopac).

The 4 min of data before the first tone of each arousal was used for all analyses.

Windows as small as 30 s can be used in studies of dynamic functional correlations during sleep (Wilson et al., 2015). Using the data immediately before the first tone ensured the functional correlations would be tightly associated with the functional state that led to the AAT without being influenced by ascending state transitions or evoked activity. Although sleep states change and phasic events occur on faster timescales than every 4 min, sleep stages are normally very stable, transitions can be said to be controlled by a bistable switch (Saper et al., 2010), and stability of sleep stage is linked to waking cognitive outcomes (e.g., Mazzoni et al., 1999). Only runs from the second night with ample, good-quality functional MRI and peripheral data were analyzed. This resulted in 77 four-min segments across the 12 subjects.

Using a nearly identical approach compared to our prior work on cerebrospinal fluid pulsations (Picchioni, Ozbay, et al., 2022), the functional MRI data were processed using AFNI Version 21.1.07 or later with its afni_proc.py function (Reynolds et al., 2024) in the following order. Outliers, polynomial trends, and two harmonics of the cardiac and respiratory signals from a modified “retrospective correction of physiological motion effects” model (Glover et al., 2000) were removed. Slice-timing correction; motion registration, regression, and censoring; anatomical skull stripping; and nonlinear alignment to the Talairach template were performed.

Nonlinear alignment to template space and anatomical skull stripping were performed with AFNI’s @SSwarper function prior to running afni_proc.py (Saad et al., 2009). The final volumetric element (voxel) size from this processing was 2.0 mm isotropic.

The “retrospective correction of physiological motion effects” modelling aims to remove fast, pulsatile artifacts in the functional MRI data; it does not remove artifacts caused by slower changes in autonomic nervous system tone. These were removed by calculating and separately regressing out two additional peripheral measures (Duyn et al., 2020). These variables were calculated from the respiratory effort and finger photoplethysmography signals as follows. To remove artifactual spikes from the data, time points exceeding four *SD* above the mean were set to the mean. Low-pass filtering was performed on the respiratory effort and photoplethysmography signals at 30 and 10 Hz, respectively. From the photoplethysmography signal, an indicator of peripheral vascular volume was created by calculating the *SD* of the signal in 3-s segments (the MRI volume repetition time); this is a measure of the excursion amplitude caused by the heart beat and is proportional to the volume of blood detected by the sensor (Allen, 2007). The filtered respiratory signal was used to derive a measure of respiratory flow rate, which previously was found to affect the functional MRI signal throughout the brain (Birn et al., 2008), albeit not significantly (Tu & Zhang, 2022). This was done by taking the derivative of the low-pass filtered signal, rectifying it, and applying a second low-pass filter at 0.13 Hz. To regress out the respiratory flow rate and peripheral vascular volume signals, two time-shifted versions were used for each. Shifts of 12 and 15 s were used for respiratory flow rate, and 0 and 3 s were used for peripheral vascular volume. These values were expected to be effective in accounting for peripheral contributions to the functional MRI signal (Ozbay et al., 2019). To reduce the effect of outliers, the regression was performed in five-minute segments by zero-padding the regressors at five-minute intervals.

Simultaneous EEG data were collected and sleep staged using standard procedures as previously described (Moehlman et al., 2019). The modal sleep stage in the four minutes prior to the first tone was assigned as the sleep stage for that arousal. If one stage did not obtain a majority, the sleep stage immediately before the first tone was used.

### Experimental Design and Statistical Analysis

#### Analysis 1

Correlation matrices were created where every pairwise correlation between predefined networks can be presented. In this context, a network is defined as a group of regions. This allowed examination of multiple relationships without the mental indigestion that would arise from using individual voxels or regions. A custom Talairach-space atlas was created by beginning with the Eickhoff-Zilles atlas (Eickhoff et al., 2005). The original atlas had 116 cortical and subcortical regions, which were created using both cytoarchitectural data and multiple task-based functional-activation maps. The cerebellar vermis regions were divided into left and right regions. The total was 124 left and right regions of interest.

These regions were manually categorized into 26 left-and right-hemisphere networks. In contrast to creating networks based solely on resting-state intrinsic correlation networks, this approach facilitated the inclusion of subcortical networks, allowed the potential analysis of both networks and regions, and had fewer assumptions because the regions were created from anatomy and activations associated with specific, ecologically relevant tasks. The manual categorizations were based on well-established networks (Menon, 2011; Raichle, 2011; Yeo et al., 2011) and reasoned categorizations by others (Andrews-Hanna et al., 2010; Mesulam, 1998; Vernet et al., 2014). For example, primary sensory and motor cortices were classed into a unimodal cortical network because functional correlations are anchored by regions serving primary sensory/motor functions at one end of a principal gradient of cortical hierarchical organization (Margulies et al., 2016). The resulting networks were labelled as (1) default-mode network posterior cingulate cortex-anterior medial prefrontal cortex core, (2) default-mode network dorsal medial prefrontal cortex subsystem, (3) default-mode network medial temporal lobe subsystem, (4) dorsal attention, (5) ventral attention, (6) executive control, (7) language, (8) salience, (9) heteromodal misc, (10) unimodal cortical, (11) thalamus, (12) basal ganglia, and (13) cerebellum. Although it is allocortex, as others have done, the hippocampus was categorized with neocortical regions in the “default-mode network medial temporal lobe subsystem” because its activity has a strong positive correlation with other regions in this subsystem (Andrews-Hanna et al., 2010). Although the amygdala is typically not included in this subsystem, it was added because others have grouped it with the hippocampus (Bastuji et al., 2024) and because it has a tight association with the functions of this subsystem.

The spatial mean of time points for the voxels in a network was calculated at each time point. This resulted in one functional MRI time series per network per run. No spatial smoothing was used. Spatial smoothing can be considered moot in this approach because voxels are averaged across a large spatial extent for each network. This took the place of spatial smoothing, provided similar statistical advantages, and provided a similar control for the residual geometric distortion that is present in all functional MRI data.

All pairwise correlations between network time series were calculated with AFNI’s 3dNetCorr (Taylor & Saad, 2013). The function was used to output the Fisher *z*-transformed correlation values. The absolute value for each transformed correlation was computed because a strong positive correlation and a strong negative correlation would both indicate the same magnitude of assumed neural communication/information exchange between two brain networks (e.g., Picchioni et al., 2014; Zalesky et al., 2010). This approach is particularly appropriate when considering that activity in all regions of the brain can be evoked with a task, including default-mode network regions (e.g., Gusnard et al., 2001; Spreng et al., 2009).

Separately for each correlation, the transformed absolute correlation values were entered into a linear mixed effects analysis whereby each subject can contribute more than one pair of observations (i.e., a cluster of observations for each subject). The variabilities in the slopes and intercepts across the clusters are modeled with linear mixed effects by including both the individual and cluster intercepts and slopes in the regression. The fixed effects were one continuous, within-subject variable (AAT). The random effects were AAT and the intercept across subjects. The criterion variable was the correlation between any two networks. In a preliminary analysis, another fixed effects variable was included: the discrete, within-subject variable of condition with approximately six levels representing arousals across the night (i.e., Arousal 1, Arousal 2, etc). This proved unnecessary because its effects were not examined, so it was removed for the final version of the analysis.

The linear mixed effects analysis was performed with the fitlme function in MATLAB R2022a or later (The Mathworks. Multiple testing correction was performed with the fdr_bh function. The effect size used was Cohen’s *f* ^2^. Its estimate was calculated by subtracting the proportion of variance explained by *R*^2^ in a reduced model without AAT from a full model with AAT. *R*^2^ was ordinary and conditional.

Correlation matrices are important but are limited because they are constrained by the specific topology of the preselected networks. Three exemplar regions were chosen to create seed-region correlation maps of the correlation with AAT: bilateral hippocampus, thalamus, and posterior cingulate cortex. The hippocampus was chosen given its relevance to sleep-dependent memory consolidation (e.g., Rasch et al., 2007), the thalamus was chosen given its relevance to sleep regulation (e.g., Steriade & McCarley, 2005), and the posterior cingulate cortex was chosen given its putative role in dreaming (e.g., Picchioni, Abdollahi, et al., 2022).

The spatial mean for the voxels in a seed region was calculated at each time point. The whole-brain data were then spatially smoothed to 4 mm full width at half maximum. The vectors representing each seed region’s spatial mean across time were then correlated with all voxels in the brain. Identical to the network approach, absolute values of the resulting Fisher *z*-transformed correlations were fed into a linear mixed effects analysis with AAT. In contrast to the network approach, AFNI’s 3dLMEr (Chen et al., 2013) was used to enter the whole-brain maps as the inputs into the linear mixed effects model.

Multiple testing correction was performed on the group-level results as follows. After the correlations with the seed regions were calculated, the residuals were computed and the total extent of spatial smoothness in the residuals for each subject was calculated using non-Gaussian filtering (Cox et al., 2017a, 2017b). This obviates the minor problem associated with assuming normality in the smoothness of neuroimaging data (Eklund et al., 2016). These smoothness estimates in the residuals were averaged across all subjects and fed into AFNI’s 3dClustSim function, which uses Monte Carlo simulations to estimate the size of false-positive clusters. The following parameters were used: a two-sided test, an individual-voxel *α* = 0.05, a cluster-size *α* = 0.05, and a clustering technique of nearest neighbor with 26 face, edge, and cornerwise neighbors. This gave a minimum cluster size of 1,226, 1,195, and 1,203 voxels for the hippocampus, thalamus, and posterior cingulate cortex, respectively. All statistical tests in this analysis were two-sided.

#### Analysis 2

In this analysis, the AAT-based model was directly compared to a model based on conventional EEG sleep stages. Separate statistics were calculated for AAT and stage. A spatial conjunction analysis was then performed to compare the significant results. The endpoint of interest was picture elements (pixels) in the correlation matrices and voxels in the brain maps. One color was assigned to pixels/voxels only significant for the model with the discrete variable of stage, another color was assigned to pixels/voxels only significant for the model with the continuous variable of AAT, and a third color was assigned to overlapping pixels/voxels.

For the network approach, two linear mixed effects models were created: one with the continuous variable of AAT and the second with a set of 4 (the 5 conventional stages − 1) dummy-coded predictor variables coding the discrete variable of stage. The type of dummy-coding used was effects coding where the reference level is coded as −1. These results from MATLAB’s fitlme function were fed directly into MATLAB’s anova function with an approach that is specific to analyzing the main effects from a linear mixed effects model (https://www.mathworks.com/help/stats/linearmixedmodel.anova.html). A very similar approach was used for the seed-region approach with AFNI’s 3dLMEr. The main effects of stage and AAT were calculated with chi-square with two degrees of freedom, whereas *F*-tests were used in the network approach using the Satterthwaite approximation for the degrees of freedom. Most other parameters were identical to the analyses from Analysis 1. One-sided tests were instead used for AAT and stage because directionality was not of interest. Calculating the significance of the overall variance explained by AAT and stage and then comparing those results spatially was the outcome of interest in this analysis.

## Statistical Conventions

Data are presented as *M* ± *SD*. Probability thresholds (*α*) were 0.05. To facilitate a better spatial interpretation, some have recommend using *α* = 0.001 as the primary threshold (i.e., before multiple testing correction) in neuroimaging analyses so that clusters of statistically significant voxels do not span multiple neuroanatomical regions (Woo et al., 2014). As an orthogonal and better solution, unthresholded results were prepared to complement thresholded results to minimize the practice of dichotomizing data (Cohen, 1994; Taylor et al., 2023).

Unthresholded statistical brain maps were masked to exclude voxels in white matter and cerebrospinal fluid. For a similar reason, effect sizes are presented to complement null hypothesis testing (Chen et al., 2017; Keppel, 1991).

## Results

### Descriptive Results

All-night functional MRI data were analyzed from 12 subjects. The mean age was 24.0 (*SD* = 3.5), and 33.3% were male. An adaptation night was used to acclimate subjects to sleeping in the scanner. Auditory tones of increasing intensity were administered randomly on both nights to determine AAT as a behavioral measure of sleep depth. Only runs from the second night with ample, good-quality functional MRI and peripheral data were analyzed, and the 4 min of functional MRI data before the first tone were extracted. This resulted in 77 four-min segments across the 12 subjects. The amount of excluded data due to motion within each of the 77 four-min segments was 12.0 ± 22.0 s. Though arousals were delivered randomly to search for new sleep states temporally unconstrained by the conventional sleep stages, it is still prudent to examine the stages present in the four-min segments. Table 1 shows the amount of each conventional sleep stage, and Figure 1 shows a standard box plot of AAT for each sleep stage. It is clear from this figure that much variability exists within each stage, so AAT is capturing variance missed by the conventional sleep stages, and this variability should provide unique information about sleep depth.

**Figure 1.**
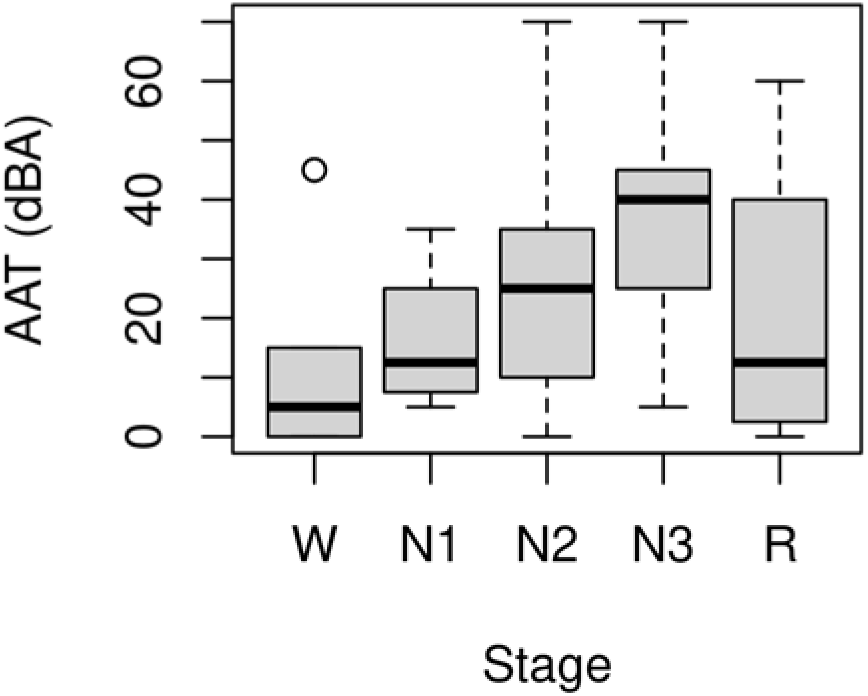
Standard box plot of auditory arousal thresholds (AAT) for each conventional sleep stage. It is clear from this figure that much variability exists within each stage, so AAT is capturing variance missed by the conventional sleep stages, and this variability should provide unique information about sleep depth. W = stage wakefulness (*n =* 16); N1 = stage nonrapid eye movement 1 sleep (*n* = 4); N2 = stage nonrapid eye movement 2 sleep (*n* = 32); N3 = stage nonrapid eye movement 3 sleep (*n* = 21); R = stage rapid eye movement sleep (*n* = 4).

**Table 1.** Descriptive Statistics of the Conventional Stages Present in the Four-min Functional MRI Segments.

| Stage | *M*<br>(s) | *SD*<br>(s) | Modal<br>Stage<br>(%) |
| --- | --- | --- | --- |
| W | 48.9 | 83.1 | 20.8 |
| N1 | 22.7 | 44.0 | 5.2 |
| N2 | 83.1 | 86.9 | 41.6 |
| N3 | 72.9 | 94.3 | 27.3 |
| R | 12.4 | 49.5 | 5.2 |

Note: Though arousals were delivered randomly to search for new sleep states temporally unconstrained by the conventional sleep stages, it is still prudent to examine the stages present in the four-min functional magnetic resonance imaging (MRI) segments. $n = 77$ . W = stage wakefulness; N1 = stage nonrapid eye movement 1 sleep; N2 = stage nonrapid eye movement 2 sleep; N3 = stage nonrapid eye movement 3 sleep; R = stage rapid eye movement sleep.

### Analysis 1

AAT-dependent correlation metrics are presented in Figures 2-4. Figure 2 displays unthresholded statistical tests, thresholded statistical tests, and effect sizes for the correlation between AAT and the correlation between pairs of networks. The most salient feature is the negative corticocortical correlations. This means that as AAT/sleep depth increased, corticocortical functional correlations decreased. Negative thalamocerebellar correlations are also clear from the thresholded matrix as were negative associations with AAT and functional correlations between the thalamus and executive control network.

**Figure 2.**
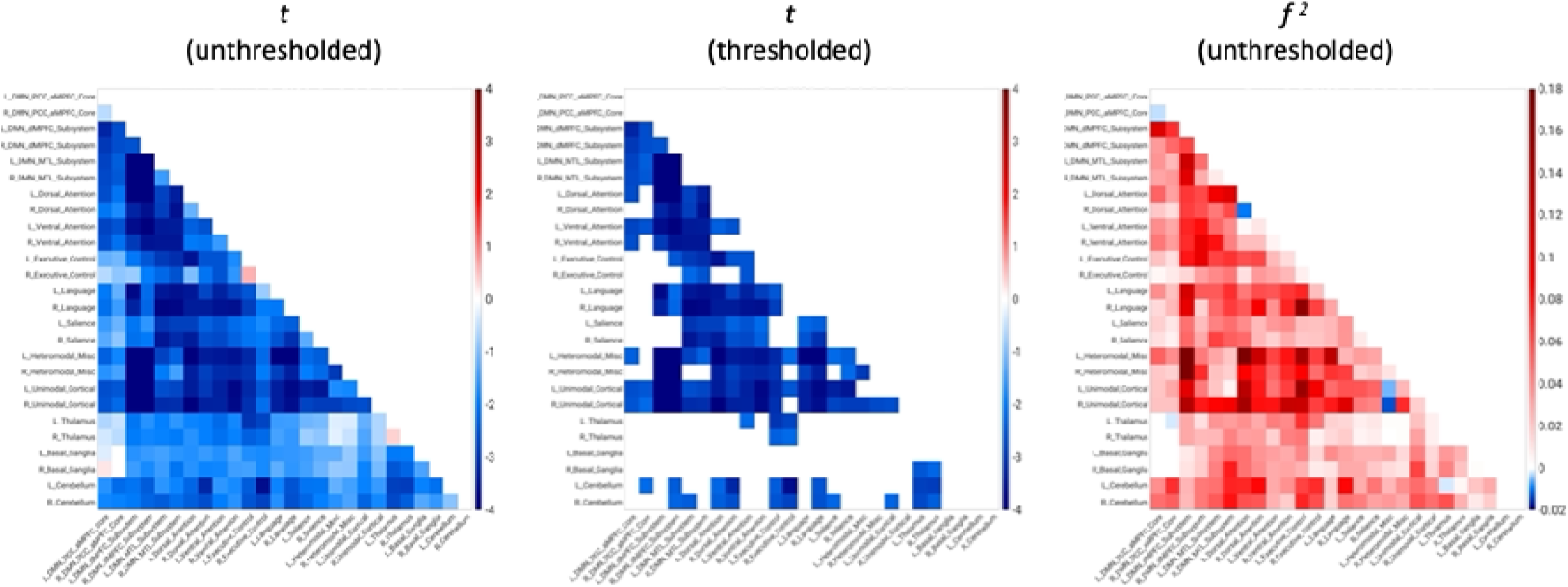
Unthresholded and thresholded linear mixed effects correlations between auditory arousal thresholds and pairs of network correlations for all arousals (*n* = 77). Thresholded results use an individual-pixel probability threshold of 0.05 plus a false-discovery rate multiple testing correction. The most salient feature is the negative corticocortical correlations. This means that as AAT/sleep depth increased, corticocortical functional correlations decreased. *t* = *t*-test. *f* ^2^ = Cohen’s *f* ^2^ effect size. DMN PCC aMPFC = default-mode network posterior cingulate cortex-anterior medial prefrontal cortex. DMN dMPFC = default-mode network dorsal medial prefrontal cortex. DMN MTL = default-mode network medial temporal lobe.

**Figure 3.**
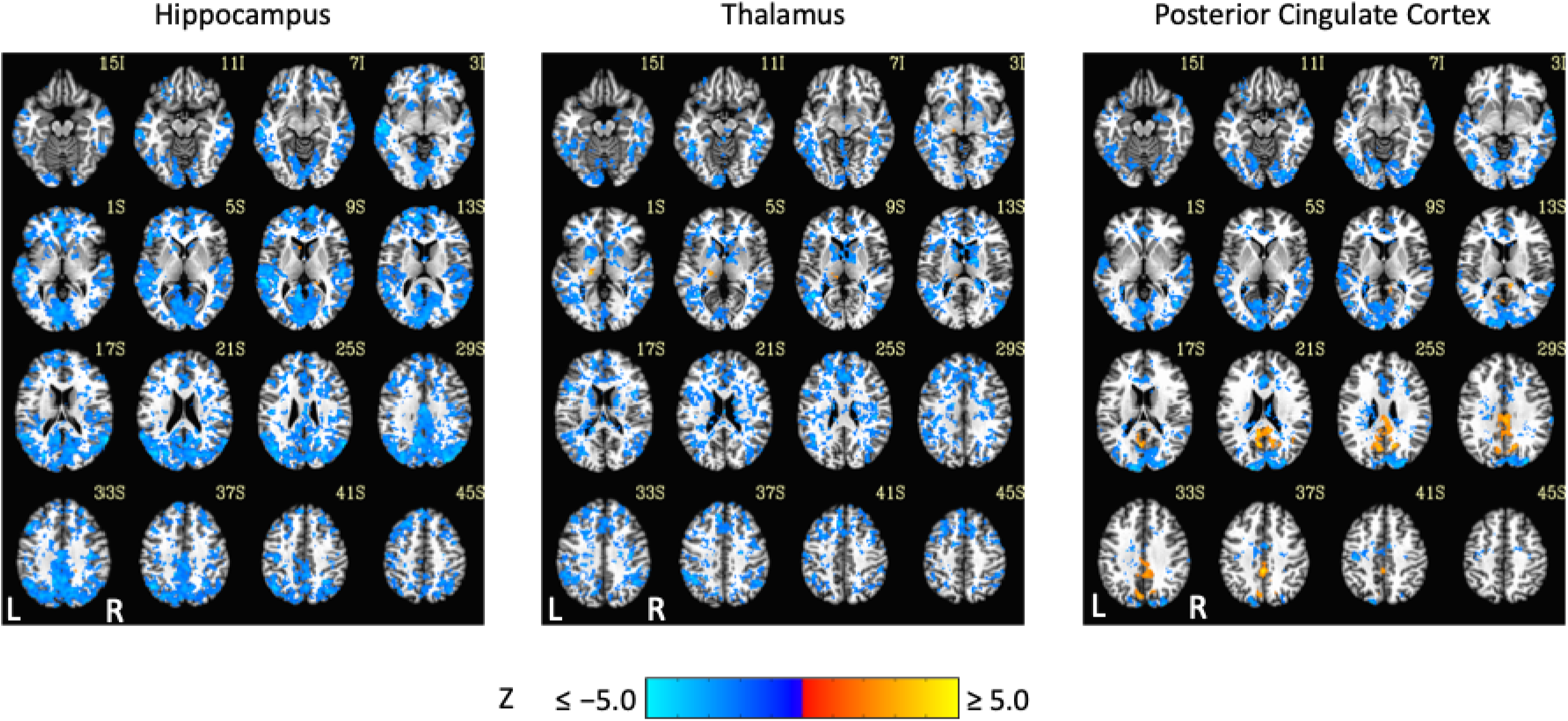
Thresholded linear mixed effects correlations between auditory arousal thresholds (AAT) and seed-region correlations for three exemplar seed regions. Decreases in corticocortical correlations observed at the network level can be observed for the posterior cingulate cortex seed region. Moreover, as AAT/sleep depth increased, thalamic correlations with the neocortex decreased. A two-sided test, an individual-voxel probability threshold of 0.05, a cluster-size probability threshold of 0.05, and a clustering technique of nearest neighbor with 26 face, edge, and cornerwise neighbors were used. This gave a minimum cluster size of 1,226, 1,195, and 1,203 voxels for the hippocampus, thalamus, and posterior cingulate cortex, respectively.

**Figure 4.**
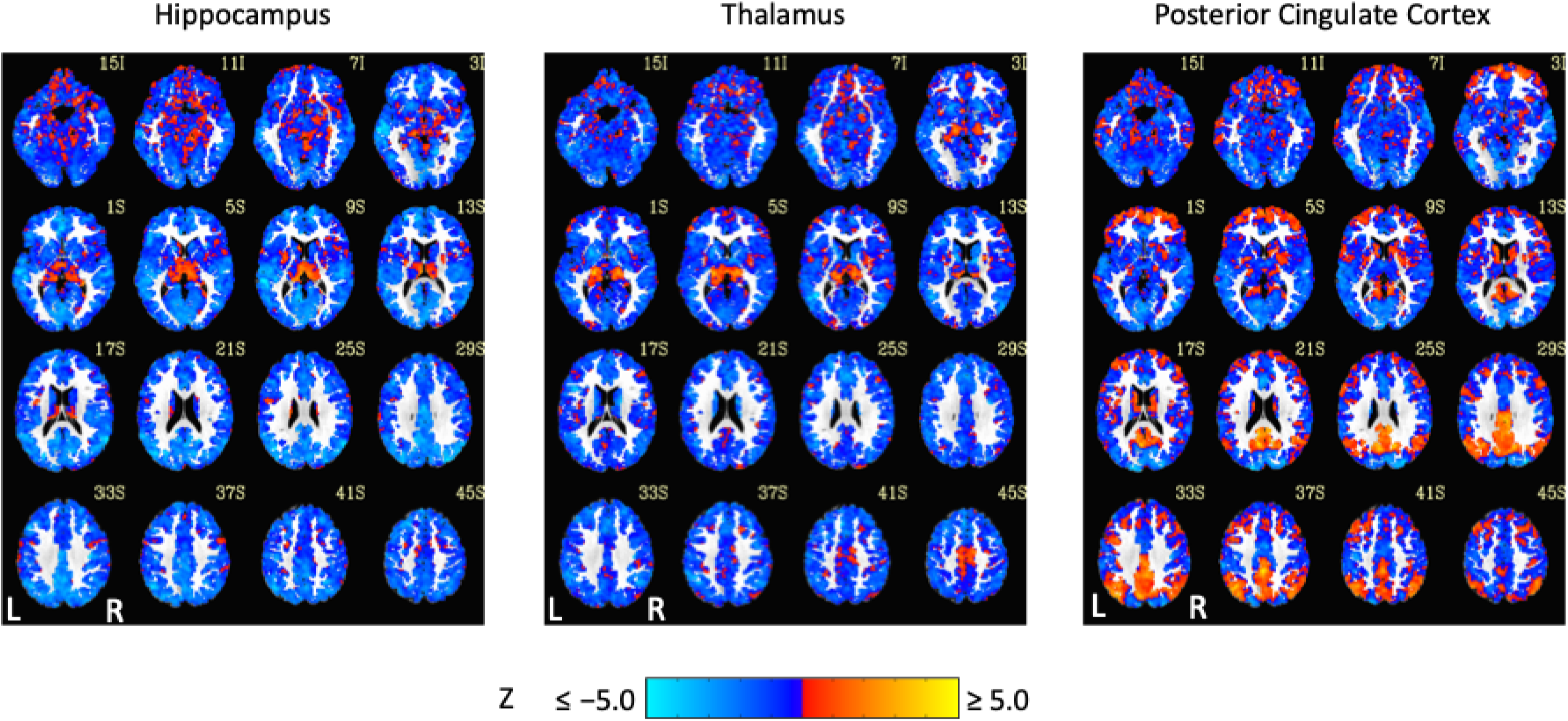
Unthresholded linear mixed effects correlations between auditory arousal thresholds and seed-region correlations for three exemplar seed regions with a white matter plus cerebrospinal fluid mask. These results were prepared to complement thresholded results to minimize the practice of dichotomizing data with arbitrarily strict probability thresholds.

Exemplar seed-based, AAT-dependent correlation metrics are shown in Figures 3 and 4. Decreases in corticocortical correlations observed at the network level can be observed for the posterior cingulate cortex seed region. For the hippocampus, similar results were found, and this is not surprising given that the hippocampus is often considered part of the default-mode network (Andrews-Hanna et al., 2010). There were some positive correlations between AAT and the relationship between the thalamus with itself. This result was seen in the unthresholded network approach, but it did not survive statistical thresholding. More broadly, as AAT/sleep depth increased, thalamic correlations with the neocortex decreased, a result that replicates prior work with a different dataset (Picchioni et al., 2014). This was seen in the thresholded network approach for the relationship between the thalamus and the executive control network.

A possible concern for this study is the effect of stage REM sleep. One of the primary goals of performing all-night functional MRI sleep studies was to obtain the full spectrum of sleep stages, conventional or undiscovered, during a night of human sleep, but to compare the current results to prior studies, the majority of which did not have any stage REM sleep, and to simplify the analysis in terms of restricting it to NREM sleep processes, the network approach was repeated while excluding the four arousals from this stage. Figure 5 recreates Figure 2 with the only difference being the exclusion of these four arousals. Aside from a few negative thalamic pixels that now passed the threshold mask, the results were very similar.

**Figure 5.**
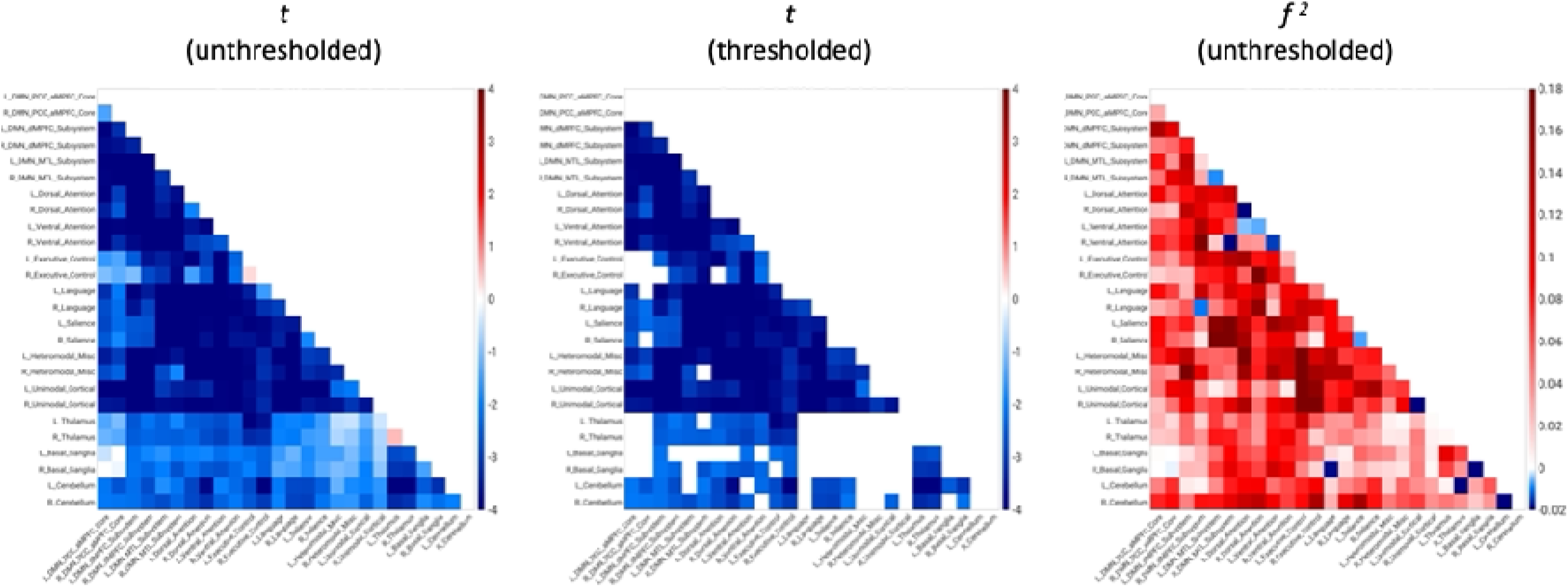
Unthresholded and thresholded linear mixed effects correlations between auditory arousal thresholds and pairs of network correlations for the arousals (*n* = 73) *not* classified as stage rapid eye movement (REM) sleep arousals. One of the primary goals of performing all-night functional MRI sleep studies was to obtain the full spectrum of sleep stages, conventional or undiscovered, during a night of human sleep, but to compare the current study to prior studies, the majority of which did not have any stage REM sleep, and to simplify the analysis in terms of restricting it to nonrapid eye movement sleep processes, the network approach was repeated while excluding the four arousals from this stage. This figure recreates Figure 2 with the only difference being the exclusion of these four arousals. Aside from a few negative thalamic pixels that now passed the threshold mask, the results were very similar.

### Analysis 2

Comparison of functional MRI correlation patterns specific to AAT versus sleep stage are presented in Figures 6-8. In the brain maps, for example, stage was uniquely related to the functional correlations in the precuneus/posterior cingulate cortex for the thalamic seed region.

**Figure 6.**
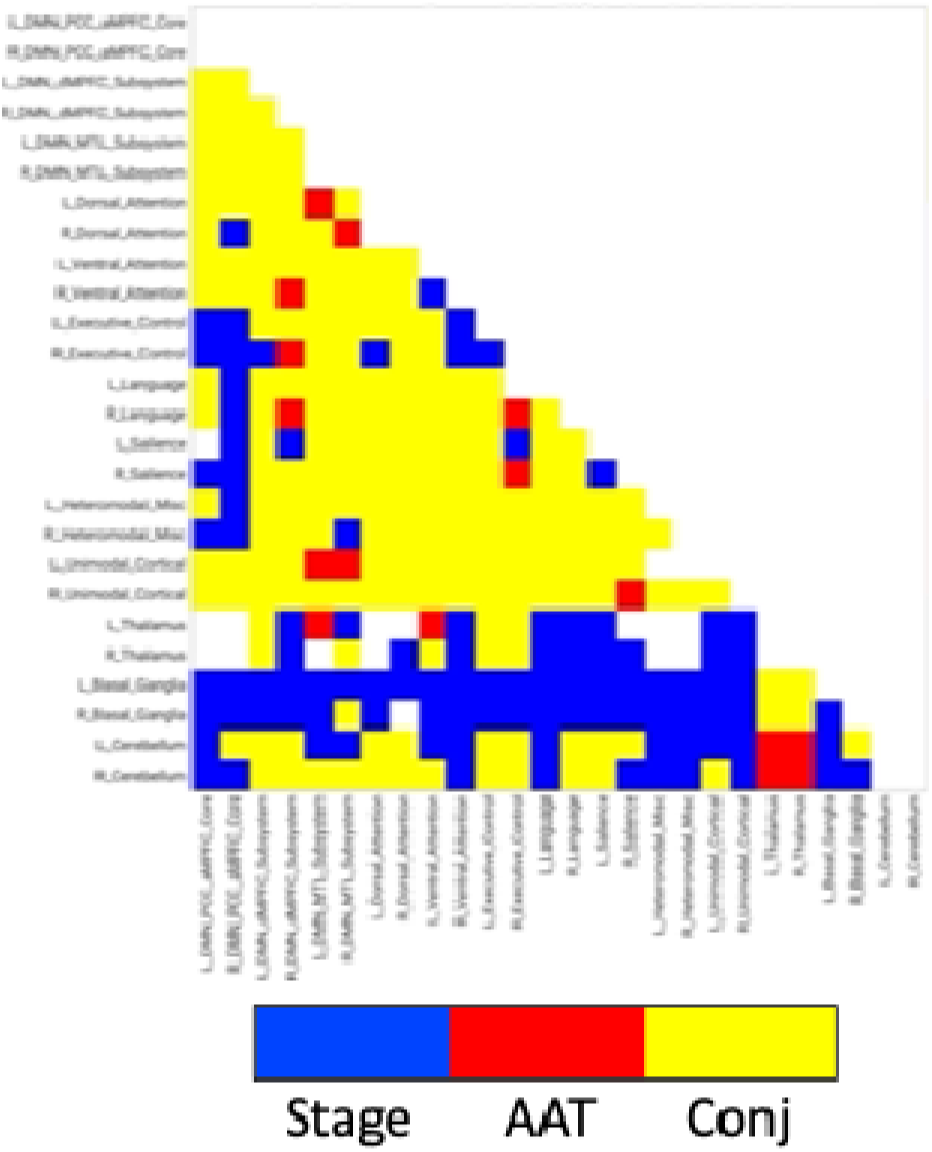
Network results of the spatial conjunction analysis of significant pixels for the continuous variable of AAT and the discrete variable of stage, represented in a separate model as a set of 4 (the 5 conventional stages − 1) dummy-coded predictor variables. Blue represents pixels only significant for the model with the discrete variable of stage, red represents pixels only significant for the model with the continuous variable of AAT, and yellow represents overlapping pixels. AAT = auditory arousal threshold. Conj = conjunction.

**Figure 7.**
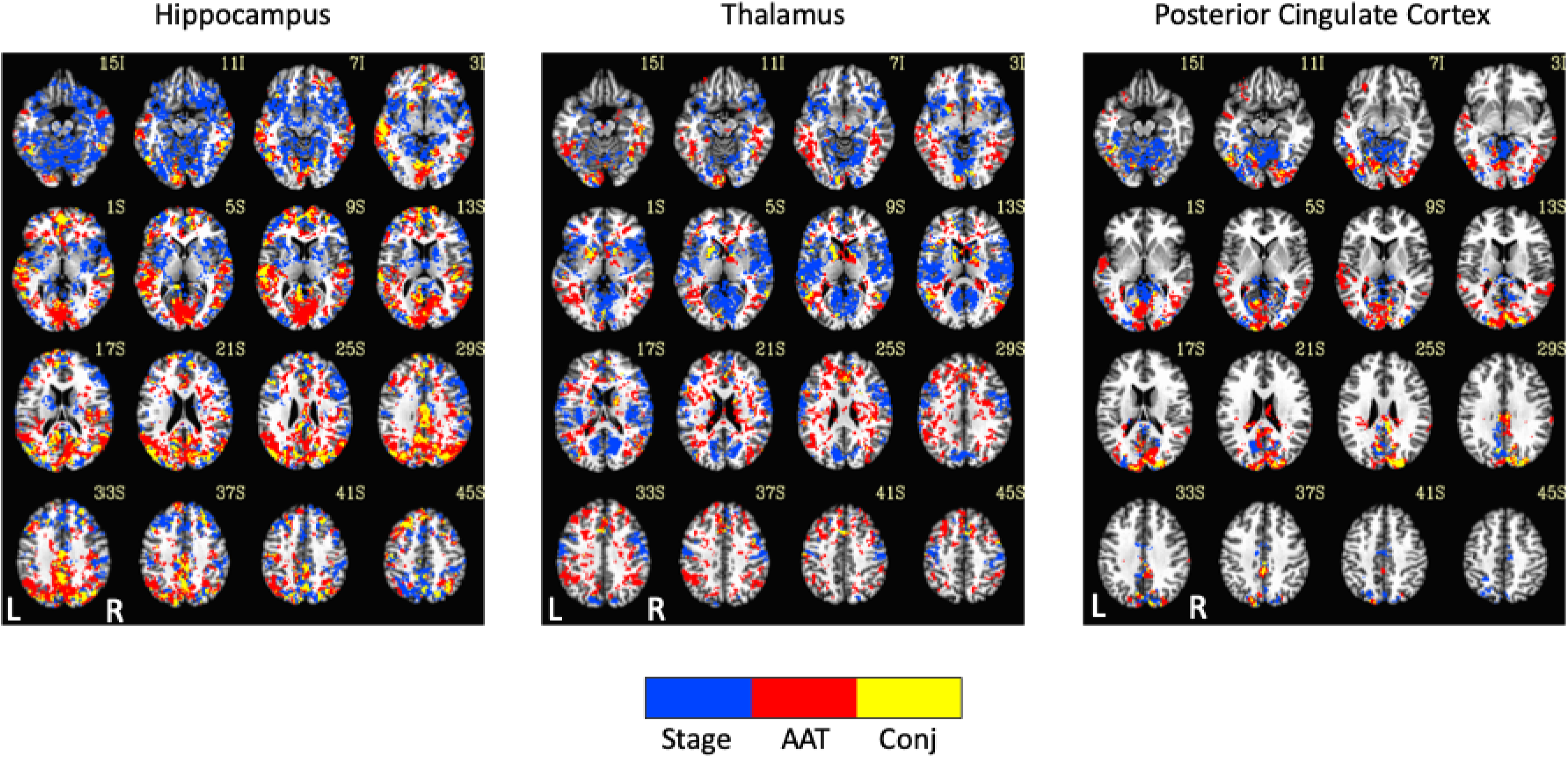
Seed-region results of the spatial conjunction analysis of significant voxels for the continuous variable of AAT and the discrete variable of stage, represented in a separate model. Blue represents voxels only significant for the model with the discrete variable of stage, red represents voxels only significant for the model with the continuous variable of AAT, and yellow represents overlapping voxels. AAT = auditory arousal threshold. Conj = conjunction.

**Figure 8.**
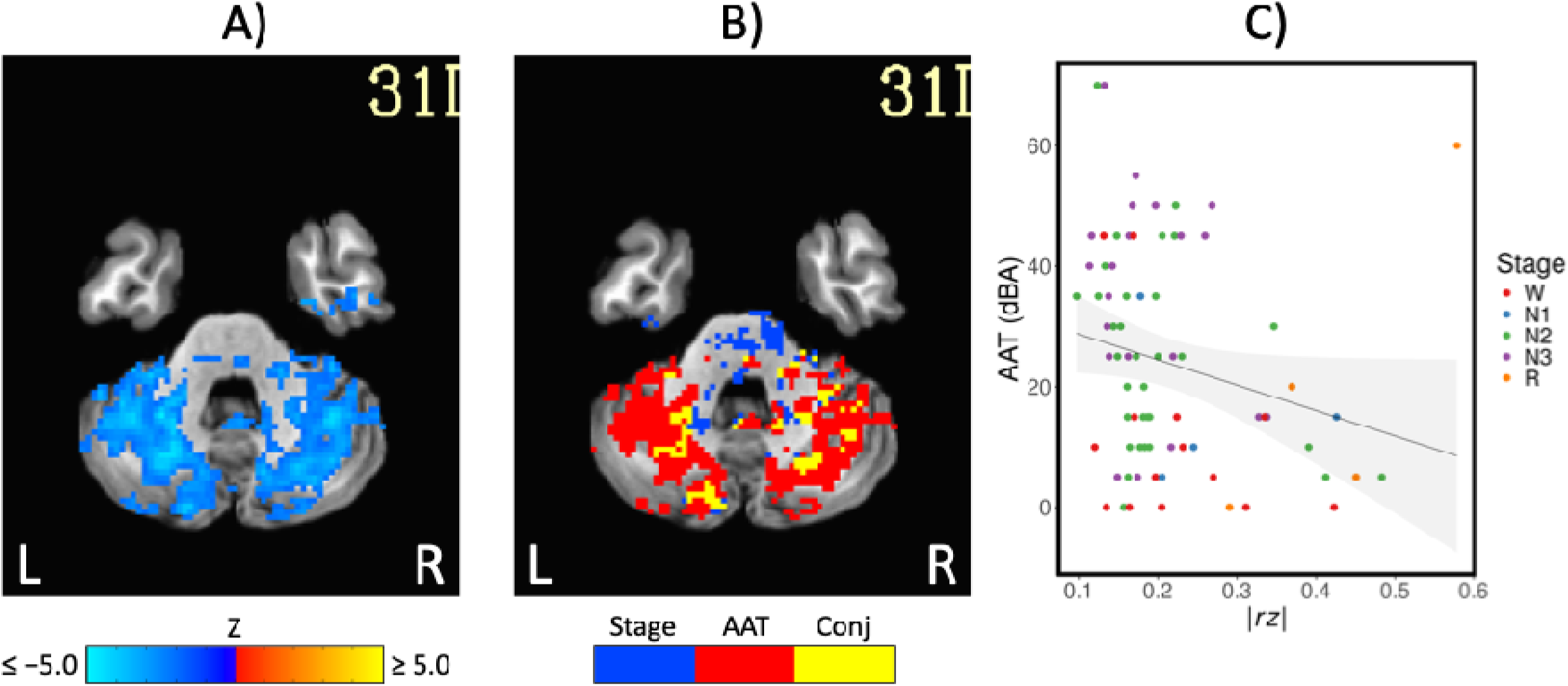
A) The negative correlations with auditory arousal threshold for the thalamic seed region. This is the same result from Figure 3 at a more-inferior slice through the cerebellum. B) The uniquely significant relationships with auditory arousal threshold for the thalamic seed region. This is the same result from Figure 7 at a more-inferior slice. C) Scatterplot of the negative relationship between absolute z-transformed functional correlations (*r z*) in a cerebellum region of interest and auditory arousal threshold (AAT) for the thalamic seed region. Conj = conjunction.

In the correlation matrix in Figure 6, relationships between basal ganglia and nearly all regions showed a unique relationship with the conventional sleep stages. Pixels with overlapping significance between AAT and sleep stage constituted most of the matrix. However, many pixels in Figure 6 and voxels in Figure 7 were uniquely significant for AAT. In the brain maps, clusters that were significant for both stage and AAT were commonly surrounded by larger clusters that were uniquely significant for AAT. In the matrix, unique relationships for AAT included thalamocerebellar correlations, the relationship between the right language network and the right “default-mode network dorsal medial prefrontal cortex subsystem,” and the relationship between thalamus and ventral attention network. Focusing on the cerebellum, correlations between bilateral thalamus and bilateral cerebellum showed a unique relationship with AAT. This is seen in the lower-right portion of Figure 6. This is also observed in the brain maps for the thalamic seed region. Figure 8A shows the same result from Figure 3 from Analysis 1 for the thalamic seed region but at a more-inferior slice through the cerebellum. The thalamocerebellar relationship with AAT is clearly seen. Figure 8B does the same for the conjunction analysis from Figure 7, and it is again clearly observed that the thalamocerebellar relationships were mostly unique for AAT. Thus, this result was highly consistent. All the points for it are given in Figure 8C. Altogether, the findings of unique functional correlation relationships with AAT in this analysis support the idea that AAT provides unique information above and beyond the conventional EEG sleep stages.

## Discussion

### Summary

The current study used all-night functional MRI sleep data and defined sleep behaviorally with AAT to characterize sleep depth better by searching for novel neural markers of sleep depth that are neuroanatomically localized and temporally unrelated to the conventional stages.

### Comparisons with Existing Literature

It is first important to note the finding of a decrease in corticocortical correlations with decreases in consciousness (i.e., increases in AAT) replicates other studies using the conventional sleep stages or similar methodology, including those that do not rely on blood flow as a proxy for neural activity. Examples include human studies of natural sleep using low-density scalp EEG (De Gennaro et al., 2001), high-density scalp EEG (Titone et al., 2024), intracranial EEG (Bastuji et al., 2021), functional MRI (Picchioni et al., 2013), and simultaneous EEG-transcranial magnetic stimulation (Massimini et al., 2005). These results have been extended to human studies of coma and anesthesia (Noirhomme et al., 2010), human research on intracerebellar-cortex correlations (Liu et al., 2023), and nonhuman animals (Olcese et al., 2016; Tu et al., 2021). This is explained by an overall reduction in brain activity (Braun et al., 1997) and/or a relative increase in offline and isolated information processing during up and down states in neocortical neurons (Destexhe et al., 2007) as opposed to the coordinated information processing between regions during goal-directed behavior in wakefulness.

Arousal threshold was uniquely related to the relationship between bilateral cerebellum and bilateral thalamus. Thalamocerebellar connectivity is just as important for understanding sleep as thalamocortical connectivity (Steriade et al., 1971). Recently, it was found that thalamocerebellar functional MRI correlations were significantly higher in wakefulness than stage N2 and N3 sleep (Liu et al., 2023), and sleep spindles, which are phasic bursts of 11-16 Hz oscillations, were found to cooccur with a directionally specific flow of information from the cerebellum to the thalamus to the motor cortex (Xu et al., 2021). Sleep spindles are the first marker of substantial increases in arousal threshold during sleep onset (Bonnet & Moore, 1982), and further support for this interesting connection between sleep spindles, the cerebellum, and sleep depth would have been left undiscovered if one had analyzed the data according to the conventional stages.

Language is a vital part of cognition; cognition is a vital part of consciousness; and consciousness is a vital part of the self. Many definitions of consciousness incorporate an awareness of self. This contextualizes another unique relationship for AAT: the relationship between the right language network and the right “default-mode network dorsal medial prefrontal cortex subsystem,” which is commonly labelled as the self-referential subsystem (Molnar-Szakacs & Uddin, 2013). One would expect both stage and AAT to be important for correlations between these networks. However, it was only significant for AAT, suggesting this type of analysis may be more sensitive in detecting sleep-depth dependent changes in functional correlations.

Animals must be able to maintain sleep in the face of disturbing stimuli to absorb its functions and, simultaneously, must be able to attend to relevant stimuli in the environment. Children can sleep through sounds equivalent to a blender or garbage disposal (e.g., Busby et al., 1994), yet hearing one’s name can penetrate the veil of sleep and cause brain and behavioral responses (Perrin et al., 1999; Voss & Harsh, 1998). This comes into play when measuring sleep depth via arousal threshold. One would expect a relationship between thalamus and ventral attention network because, like the thalamus acts as a gatekeeper generally, ventral attention network acts as a bottom-up attentional gatekeeper for salient stimuli, but this relationship was only significant for AAT. Again, this may indicate that a superior sensitivity of AAT-based analysis allowed it to detect new markers of sleep-depth regulation.

The use of arousal threshold in the context of the behavioral definition of sleep is not the only way to characterize sleep with functional neuroimaging. Others have searched for novel spatial brain states during sleep using a Hidden Markov Model (Stevner et al., 2019; Yang et al., 2024). This model is a data reduction technique and characterizes unique brain states based on time series and infers the transitions between those states. Although the investigators used the conventional stages to validate the results, the models were blind to stage when the states and transition probabilities were determined. This method identified 21 recurring brain states (Yang et al., 2024) and their transition probabilities, beyond PSG-defined sleep stages. This synergizes with the current work because important results are always found to be reproducible across different approaches to the same problem.

### Weaknesses and Strengths of the Current Study

It is possible that blood flow was affected by other factors such as autonomic nervous system tone (Duyn et al., 2020). This was addressed in the current study by removing the variance associated with measures of autonomic tone, but these measures are only approximations.

It cannot be said that the sample size in this study was small. It can be said that the necessary sample size to detect the effect of interest was not known because an *a priori* statistical power analysis was not performed. With the data from larger studies, other analyses can be performed, such as including both stage and AAT in the same model to measure the unique variance explained by AAT. This would complement the spatial conjunction analysis reported in the current study.

Common to all studies that use linear correlations, curvilinear relationships between AAT and functional correlations could have been missed. Future studies should include alternative techniques such as polynomial regression, entropy (Watanabe et al., 2014), information-theory measures (Olcese et al., 2016), or distance correlations (Bolt et al., 2024). However, each of these measures have limitations of their own, so they must be considered as complementary to their linear equivalents.

With very few exceptions (Picchioni, Ozbay, et al., 2022; Yang et al., 2024), analyses performed on all-night functional MRI data do not exist. Moreover, most similar studies used sleep deprivation, scanned for short durations, and/or began scanning at times that would cause circadian misalignment. These methods and the amounts of sleep obtained with them restrict the generalizability of the results, limiting it to diurnal sleep, a limited number of sleep stages (conventional or otherwise), the recovery sleep that occurs after sleep deprivation, the specific stages that occur at different parts of the night, and/or a single sleep cycle. The current analyses were performed on data without these limitations. By using arousal thresholds, the current work also avoided the circularity associated with studying brain activity/functional correlations associated with the conventional stages.

### Further Research

Searching for sleep states that are independent from the conventional stages (Nir & Tononi, 2010) and associated with dreaming would also be important. This could be done with an analysis that is like the current study’s but with dream recall rather than AAT in the model. The current study focused on the four minutes before the first tone to ensure the functional correlations would be tightly associated with the brain state that led to the arousal threshold.

Further research and analyses should capitalize on the full amount of data, for example, to search for novel states in a data driven way across the entire night. In addition to data reduction techniques that consider the entirety of the data at once, a sliding window could be used to match a template of a pattern-of-interest across the night. This has been successfully employed during sleep onset (Chang et al., 2016) but could be employed across a night of sleep.

A personalized network is a network that resembles the group average but displays differences along its borders for one subject, and these differences can have clinically significant outcomes (Fair, 2024) and reliably correlate with differences in cognition between individuals and within individuals across development (e.g., Cui et al., 2020). The canonical resting-state networks found during wakefulness have been spatially correlated with networks from sleep (Houldin et al., 2019). The idea was that networks with low spatial correlations would represent novel resting-state networks that are unique to sleep. No such networks were found, but low-correlation networks consisting of smaller constituent regions of the canonical networks existed. Perhaps this change in spatial extent is linked to the functions of sleep such as memory representations that are being refined and optimized for more efficient storage (Tononi & Cirelli, 2014). Searching for neural patterns relevant for memory consolidation during sleep would generally be important because the conventional EEG sleep stages were formed without regard to sleep function.

## Conclusion

Because sleep was defined with arousal threshold independently from the conventional sleep stages, the current study was a novel approach to studying the neuroanatomical correlates of sleep with high spatial resolution. This resulted in discoveries of novel neural markers of sleep depth that are more neuroanatomically localized and temporally unrelated to the conventional stages. This represents a single step towards a neuroanatomically localized sleep scoring system and expands our understanding of sleep and its functions beyond the constraints imposed by PSG-defined stages.

## Acknowledgements

The authors would like to acknowledge the following people and resources: Guttman, S., Harbison, S., Koretsky, A., Moehlman, T., National Institutes of Health High Performance Computing Core Biowulf Cluster, Picchioni, I., Newman, S., and Wu, T.

## Funding

This work was supported by funding from the Intramural Research Program of the National Institute of Neurological Disorders and Stroke (Project: ZIA NS003027; Crossref Funder ID: 100000065) and the National Institute of Mental Health (Project: ZIC MH002888; Crossref Funder ID: 100000025).

## Notes

These results have been posted to a preprint archive at https://www.biorxiv.org/content/10.1101/2024.08.09.607376. Raw data are available at https://doi.org/10.18112/openneuro.ds005127.v1.0.2. The afni_proc.py command for one run for one subject is available at https://github.com/dantepicchioni/sleep1_afni_proc.py. The custom atlas is available at https://osf.io/uqrab/?view_only=2ae81bc515434bcc91901dab12c2b423.

## Conflict of interest statement

none declared. The National Institutes of Health Institutional Review Board Protocol Number is 2016-N-0031, and the ClinicalTrials.gov Identifier is NCT02629107. Author contributions: designed research (DP, JAdZ, PvG, JHD), performed research (DP, JAdZ, PvG, MGCF), analyzed data (DP, FNY, JAdZ, YW, HM, PSÖ, GC, PAT, NL, MGCF, CC, JL, PvG, JHD), wrote the first draft of the paper (DP), and wrote the paper (DP, FNY, JAdZ, YW, HM, PSÖ, GC, PAT, NL, MGCF, CC, JL, PvG, JHD).

## Notes

### Competing Interest Statement

The authors have declared no competing interest.

### Summary of Updates

Some minor details about the method were added.

## References

Allen, J. (2007). Photoplethysmography and its application in clinical physiological measurement. Physiological Measurement, 28(3), R1–39. 10.1088/0967-3334/28/3/R01

Andrews-Hanna, J. R., Reidler, J. S., Sepulcre, J., Poulin, R., & Buckner, R. L. (2010). Functional-anatomic fractionation of the brain’s default network. Neuron, 65(4), 550–562. 10.1016/j.neuron.2010.02.005

Bastien, C. H., Vallieres, A., & Morin, C. M. (2001). Validation of the Insomnia Severity Index as an outcome measure for insomnia research. Sleep Medicine, 2(4), 297–307. 10.1016/s1389-9457(00)00065-4

Bastuji, H., Cadic-Melchior, A., Magnin, M., & Garcia-Larrea, L. (2021). Intracortical functional connectivity predicts arousal to noxious stimuli during sleep in humans. Journal of Neuroscience, 41(23), 5115–5123. 10.1523/jneurosci.2935-20.2021

Bastuji, H., Daoud, M., Magnin, M., & Garcia-Larrea, L. (2024). REM sleep remains paradoxical: Sub-states determined by thalamo-cortical and cortico-cortical functional connectivity. Journal of Physiology, 602(20), 5269–5287. 10.1113/JP286074

Birn, R. M., Smith, M. A., Jones, T. B., & Bandettini, P. A. (2008). The respiration response function: The temporal dynamics of fMRI signal fluctuations related to changes in respiration. Neuroimage, 40(2), 644–654. 10.1016/j.neuroimage.2007.11.059

Blake, H., & Gerard, R. W. (1937). Brain potentials during sleep. American Journal of Physiology, 119(4), 692–703. 10.1152/ajplegacy.1937.119.4.692

Blumberg, M. S., Karlsson, K. A., Seelke, A. M., & Mohns, E. J. (2005). The ontogeny of mammalian sleep: A response to Frank and Heller (2003). Journal of Sleep Research, 14, 91–98. 10.1111/j.1365-2869.2004.00430_1.x

Bolt, T., Wang, S., Nomi, J. S., Setton, R., Gold, B. P., Frederick, B. d., Yeo, B. T. T., Chen, J. J., Picchioni, D., Spreng, R. N., Keilholz, S. D., Uddin, L. Q., & Chang, C. (2024). Widespread neural and autonomic system synchrony across the brain-body axis. bioRxiv. 10.1101/2023.01.19.524818

Bonnet, M. H., & Moore, S. E. (1982). The threshold of sleep: Perception of sleep as a function of time asleep and auditory threshold. Sleep, 5(3), 267–276. 10.1093/sleep/5.3.267

Brandenberger, G., Ehrhart, J., & Buchheit, M. (2005). Sleep stage 2: An electroencephalographic, autonomic, and hormonal duality. Sleep, 28(12), 1535–1540. 10.1093/sleep/28.12.1535

Braun, A. R., Balkin, T. J., Wesenten, N. J., Carson, R. E., Varga, M., Baldwin, P., Selbie, S., Belenky, G., & Herscovitch, P. (1997). Regional cerebral blood flow throughout the sleep-wake cycle. An H2(15)O PET study. Brain, 120 ( Pt 7), 1173–1197. 10.1093/brain/120.7.1173

Bueno-Junior, L. S., Ruckstuhl, M. S., Lim, M. M., & Watson, B. O. (2023). The temporal structure of REM sleep shows minute-scale fluctuations across brain and body in mice and humans. Proceedings of the National Academy of Sciences of the United States of America, 120(18), e2213438120. 10.1073/pnas.2213438120

Busby, K. A., Mercier, L., & Pivik, R. T. (1994). Ontogenetic variations in auditory arousal threshold during sleep. Psychophysiology, 31(2), 182–188. 10.1111/j.1469-8986.1994.tb01038.x

Chang, C., Leopold, D. A., Schölvinck, M. L., Mandelkow, H., Picchioni, D., Liu, X., Ye, F. Q., Turchi, J. N., & Duyn, J. H. (2016). Tracking brain arousal fluctuations with fMRI. Proceedings of the National Academy of Sciences of the United States of America, 113(16), 4518–4523. 10.1073/pnas.1520613113

Chen, G., Saad, Z. S., Britton, J. C., Pine, D. S., & Cox, R. W. (2013). Linear mixed-effects modeling approach to FMRI group analysis. Neuroimage, 73, 176–190. 10.1016/j.neuroimage.2013.01.047

Chen, G., Taylor, P. A., & Cox, R. W. (2017). Is the statistic value all we should care about in neuroimaging? Neuroimage, 147, 952–959. 10.1016/j.neuroimage.2016.09.066

Cohen, J. (1994). The earth is round (p <.05). American Psychologist, 49(12), 997–1003. 10.1037/0003-066x.49.12.997

Cox, R. W., Chen, G., Glen, D. R., Reynolds, R. C., & Taylor, P. A. (2017a). fMRI clustering and false-positive rates. Proceedings of the National Academy of Sciences of the United States of America, 114(17), E3370–E3371. 10.1073/pnas.1614961114

Cox, R. W., Chen, G., Glen, D. R., Reynolds, R. C., & Taylor, P. A. (2017b). fMRI clustering in AFNI: False-positive rates redux. Brain Connectivity, 7(3), 152–171. 10.1089/brain.2016.0475

Cui, Z., Li, H., Xia, C. H., Larsen, B., Adebimpe, A., Baum, G. L., Cieslak, M., Gur, R. E., Gur, R. C., Moore, T. M., Oathes, D. J., Alexander-Bloch, A. F., Raznahan, A., Roalf, D. R., Shinohara, R. T., Wolf, D. H., Davatzikos, C., Bassett, D. S., Fair, D. A., Fan, Y., & Satterthwaite, T. D. (2020). Individual variation in functional topography of association networks in youth. Neuron, 106(2), 340–353 e348. 10.1016/j.neuron.2020.01.029

Datta, S. (2000). Avoidance task training potentiates phasic pontine-wave density in the rat: A mechanism for sleep-dependent plasticity. Journal of Neuroscience, 20(22), 8607–8613. 10.1523/jneurosci.20-22-08607.2000

De Gennaro, L., Ferrara, M., Curcio, G., & Cristiani, R. (2001). Antero-posterior EEG changes during the wakefulness-sleep transition. Clinical Neurophysiology, 112(10), 1901–1911. 10.1016/s1388-2457(01)00649-6

Decat, N., Walter, J., Koh, Z. H., Sribanditmongkol, P., Fulcher, B. D., Windt, J. M., Andrillon, T., & Tsuchiya, N. (2022). Beyond traditional sleep scoring: Massive feature extraction and data-driven clustering of sleep time series. Sleep Medicine, 98, 39–52. 10.1016/j.sleep.2022.06.013

Destexhe, A., Hughes, S. W., Rudolph, M., & Crunelli, V. (2007). Are corticothalamic’up’ states fragments of wakefulness? Trends in Neurosciences, 30(7), 334–342. 10.1016/j.tins.2007.04.006

Dong, Y., Li, J., Zhou, M., Du, Y., & Liu, D. (2022). Cortical regulation of two-stage rapid eye movement sleep. Nature Neuroscience, 25(12), 1675–1682. 10.1038/s41593-022-01195-2

Duyn, J. H., Ozbay, P. S., Chang, C., & Picchioni, D. (2020). Physiological changes in sleep that affect fMRI inference. Current Opinion in Behavioral Sciences, 33, 42–50. 10.1016/j.cobeha.2019.12.007

Eickhoff, S. B., Stephan, K. E., Mohlberg, H., Grefkes, C., Fink, G. R., Amunts, K., & Zilles, K. (2005). A new SPM toolbox for combining probabilistic cytoarchitectonic maps and functional imaging data. Neuroimage, 25(4), 1325–1335. 10.1016/j.neuroimage.2004.12.034

Eklund, A., Nichols, T. E., & Knutsson, H. (2016). Cluster failure: Why fMRI inferences for spatial extent have inflated false-positive rates. Proceedings of the National Academy of Sciences of the United States of America, 113(28), 7900–7905. 10.1073/pnas.1602413113

Ermis, U., Krakow, K., & Voss, U. (2010). Arousal thresholds during human tonic and phasic REM sleep. Journal of Sleep Research, 19(3), 400–406. 10.1111/j.1365-2869.2010.00831.x

Fair, D. A. (2024). Liftoff: Neuropsychiatry with functional MRI comes of age [Invited lecture]. Society for Neuroscience conference, Chicago, USA.

Flanigan, W. F., Jr., Wilcox, R. H., & Rechtschaffen, A. (1973). The EEG and behavioral continuum of the crocodilian, Caiman sclerops. Electroencephalography and Clinical Neurophysiology, 34, 521–538.

Glover, G. H., Li, T. Q., & Ress, D. (2000). Image-based method for retrospective correction of physiological motion effects in fMRI: RETROICOR. Magnetic Resonance in Medicine, 44(1), 162–167.

Guo, Y., Chen, Y., Shao, Y., Hu, S., Zou, G., Chen, J., Li, Y., Gao, X., Liu, J., Yao, P., Zhou, S., Xu, J., Gao, J. H., Zou, Q., & Sun, H. (2023). Thalamic network under wakefulness after sleep onset and its coupling with daytime fatigue in insomnia disorder: An EEG-fMRI study. Journal of Affective Disorders, 334, 92–99. 10.1016/j.jad.2023.04.100

Gusnard, D. A., Akbudak, E., Shulman, G. L., & Raichle, M. E. (2001). Medial prefrontal cortex and self-referential mental activity: Relation to a default mode of brain function. Proceedings of the National Academy of Sciences of the United States of America, 98(7), 4259–4264. 10.1073/pnas.071043098

Hendricks, J. C., Sehgal, A., & Pack, A. I. (2000). The need for a simple animal model to understand sleep. Progress in Neurobiology, 61, 339–351.

Himanen, S. L., & Hasan, J. (2000). Limitations of Rechtschaffen and Kales. Sleep Medicine Reviews, 4(2), 149–167. 10.1053/smrv.1999.0086

Houldin, E., Fang, Z., Ray, L. B., Owen, A. M., & Fogel, S. M. (2019). Toward a complete taxonomy of resting state networks across wakefulness and sleep: An assessment of spatially distinct resting state networks using independent component analysis. Sleep, 42(3). 10.1093/sleep/zsy235

Jang, R. S., Ciliberti, D., Mankin, E. A., & Poe, G. R. (2022). Recurrent hippocampo-neocortical sleep-state divergence in humans. Proceedings of the National Academy of Sciences of the United States of America, 119(44), e2123427119. 10.1073/pnas.2123427119

Kay, D. B., Karim, H. T., Soehner, A. M., Hasler, B. P., James, J. A., Germain, A., Hall, M. H., Franzen, P. L., Price, J. C., Nofzinger, E. A., & Buysse, D. J. (2017). Subjective-objective sleep discrepancy is associated with alterations in regional glucose metabolism in patients with insomnia and good sleeper controls. Sleep, 40(11). 10.1093/sleep/zsx155

Keppel, G. (1991). Design and analysis: A researcher’s handbook (3rd ed.). Prentice Hall.

Kleitman, M. (1963). Sleep and wakefulness (Revised and enlarged ed.). University Press.

Liu, J., Zou, G., Xu, J., Zhou, S., Qin, L., Sun, H., Zou, Q., & Gao, J. H. (2023). State-dependent and region-specific alterations of cerebellar connectivity across stable human wakefulness and NREM sleep states. Neuroimage, 266, 119823. 10.1016/j.neuroimage.2022.119823

Maquet, P. (2000). Functional neuroimaging of normal human sleep by positron emission tomography. Journal of Sleep Research, 9(3), 207–231. 10.1046/j.1365-2869.2000.00214.x

Maquet, P., Degueldre, C., Delfiore, G., Aerts, J., Peters, J. M., Luxen, A., & Franck, G. (1997). Functional neuroanatomy of human slow wave sleep. Journal of Neuroscience, 17(8), 2807–2812. 10.1523/jneurosci.17-08-02807.1997

Margulies, D. S., Ghosh, S. S., Goulas, A., Falkiewicz, M., Huntenburg, J. M., Langs, G., Bezgin, G., Eickhoff, S. B., Castellanos, F. X., Petrides, M., Jefferies, E., & Smallwood, J. (2016). Situating the default-mode network along a principal gradient of macroscale cortical organization. Proceedings of the National Academy of Sciences of the United States of America, 113(44), 12574–12579. 10.1073/pnas.1608282113

Massimini, M., Ferrarelli, F., Huber, R., Esser, S. K., Singh, H., & Tononi, G. (2005). Breakdown of cortical effective connectivity during sleep. Science, 309(5744), 2228–2232. 10.1126/science.1117256

Mazzoni, G., Gori, S., Formicola, G., Gneri, C., Massetani, R., Murri, L., & Salzarulo, P. (1999). Word recall correlates with sleep cycles in elderly subjects. Journal of Sleep Research, 8(3), 185–188. 10.1046/j.1365-2869.1999.00154.x

Menon, V. (2011). Large-scale brain networks and psychopathology: A unifying triple network model. Trends in Cognitive Sciences, 15(10), 483–506. 10.1016/j.tics.2011.08.003

Mesulam, M. M. (1998). From sensation to cognition. Brain, 121 (*Pt 6*), 1013–1052. 10.1093/brain/121.6.1013

Moehlman, T. M., de Zwart, J. A., Chappel-Farley, M. G., Liu, X., McClain, I. B., Chang, C., Mandelkow, H., Ozbay, P. S., Johnson, N. L., Bieber, R. E., Fernandez, K. A., King, K. A., Zalewski, C. K., Brewer, C. C., van Gelderen, P., Duyn, J. H., & Picchioni, D. (2019). All-night functional magnetic resonance imaging sleep studies. Journal of Neuroscience Methods, 316, 83–98. 10.1016/j.jneumeth.2018.09.019

Molnar-Szakacs, I., & Uddin, L. Q. (2013). Self-processing and the default mode network: Interactions with the mirror neuron system. Frontiers in Human Neuroscience, 7, 571. 10.3389/fnhum.2013.00571

Nir, Y., & Tononi, G. (2010). Dreaming and the brain: From phenomenology to neurophysiology. Trends in Cognitive Sciences, 14(2), 88–100. 10.1016/j.tics.2009.12.001

Noirhomme, Q., Soddu, A., Lehembre, R., Vanhaudenhuyse, A., Boveroux, P., Boly, M., & Laureys, S. (2010). Brain connectivity in pathological and pharmacological coma. Frontiers in Systems Neuroscience, 4, 160. 10.3389/fnsys.2010.00160

Olcese, U., Bos, J. J., Vinck, M., Lankelma, J. V., van Mourik-Donga, L. B., Schlumm, F., & Pennartz, C. M. (2016). Spike-based functional connectivity in cerebral cortex and hippocampus: Loss of global connectivity is coupled to preservation of local connectivity during non-REM sleep. Journal of Neuroscience, 36(29), 7676–7692. 10.1523/jneurosci.4201-15.2016

Ozbay, P. S., Chang, C., Picchioni, D., Mandelkow, H., Chappel-Farley, M. G., van Gelderen, P., de Zwart, J. A., & Duyn, J. (2019). Sympathetic activity contributes to the fMRI signal. Communications Biology, 2, 421. 10.1038/s42003-019-0659-0

Perrin, F., Garcia-Larrea, L., Mauguiere, F., & Bastuji, H. (1999). A differential brain response to the subject’s own name persists during sleep. Clinical Neurophysiology, 110(12), 2153–2164. 10.1016/s1388-2457(99)00177-7

Picchioni, D., Abdollahi, S., Cost, V. E., Gura, H., & Pace-Schott, E. F. (2022). The neurobiology of dreaming. In M. H. Kryger, T. Roth, C. A. Goldstein, & W. C. Dement (Eds.), Principles and practice of sleep medicine (7th ed., pp. 569–578). Elsevier.

Picchioni, D., Duyn, J. H., & Horovitz, S. G. (2013). Sleep and the functional connectome. Neuroimage, 80, 387–396. 10.1016/j.neuroimage.2013.05.067

Picchioni, D., Fukunaga, M., Carr, W. S., Braun, A. R., Balkin, T. J., Duyn, J. H., & Horovitz, S. G. (2008). fMRI differences between early and late stage-1 sleep. Neuroscience Letters, 441(1), 81–85. 10.1016/j.neulet.2008.06.010

Picchioni, D., Ozbay, P. S., Mandelkow, H., de Zwart, J. A., Wang, Y., van Gelderen, P., & Duyn, J. H. (2022). Autonomic arousals contribute to brain fluid pulsations during sleep. Neuroimage, 249, 118888. 10.1016/j.neuroimage.2022.118888

Picchioni, D., Pixa, M. L., Fukunaga, M., Carr, W. S., Horovitz, S. G., Braun, A. R., & Duyn, J. H. (2014). Decreased connectivity between the thalamus and the neocortex during human nonrapid eye movement sleep. Sleep, 37(2), 387–397. 10.5665/sleep.3422

Piéron, H. (1913). Le problème physiologique du sommeil. Masson.

Price, L. J., & Kremen, I. (1980). Variations in behavioral response threshold within the REM period of human sleep. Psychophysiology, 17(2), 133–140. 10.1111/j.1469-8986.1980.tb00125.x

Raichle, M. E. (2011). The restless brain. Brain Connectivity, 1(1), 3–12. 10.1089/brain.2011.0019

Rasch, B., Buchel, C., Gais, S., & Born, J. (2007). Odor cues during slow-wave sleep prompt declarative memory consolidation. Science, 315(5817), 1426–1429. 10.1126/science.1138581

Reynolds, R. C., Glen, D. R., Chen, G., Saad, Z. S., Cox, R. W., & Taylor, P. A. (2024). Processing, evaluating and understanding FMRI data with afni_proc.py. arxiv. 10.48550/arXiv.2406.05248

Saad, Z. S., Glen, D. R., Chen, G., Beauchamp, M. S., Desai, R., & Cox, R. W. (2009). A new method for improving functional-to-structural MRI alignment using local Pearson correlation. Neuroimage, 44(3), 839–848. 10.1016/j.neuroimage.2008.09.037

Saper, C. B., Fuller, P. M., Pedersen, N. P., Lu, J., & Scammell, T. E. (2010). Sleep state switching. Neuron, 68(6), 1023–1042. 10.1016/j.neuron.2010.11.032

Siclari, F., Baird, B., Perogamvros, L., Bernardi, G., LaRocque, J. J., Riedner, B., Boly, M., Postle, B. R., & Tononi, G. (2017). The neural correlates of dreaming. Nature Neuroscience, 20(6), 872–878. 10.1038/nn.4545

Spreng, R. N., Mar, R. A., & Kim, A. S. (2009). The common neural basis of autobiographical memory, prospection, navigation, theory of mind, and the default mode: A quantitative meta-analysis. Journal of Cognitive Neuroscience, 21(3), 489–510. 10.1162/jocn.2008.21029

Steriade, M., Apostol, V., & Oakson, G. (1971). Clustered firing in the cerebello-thalamic pathway during synchronized sleep. Brain Research, 26(2), 425–432. 10.1016/S0006-8993(71)80020-3

Steriade, M., & McCarley, R. W. (2005). Brain control of wakefulness and sleep (2nd ed.). Plenum Press.

Stevner, A. B. A., Vidaurre, D., Cabral, J., Rapuano, K., Nielsen, S. F. V., Tagliazucchi, E., Laufs, H., Vuust, P., Deco, G., Woolrich, M. W., Van Someren, E., & Kringelbach, M. L. (2019). Discovery of key whole-brain transitions and dynamics during human wakefulness and non-REM sleep. Nature Communications, 10(1), 1035. 10.1038/s41467-019-08934-3

Tarokh, L., Picchioni, D., Sasaki, Y., & Achermann, P. (in press). Advanced methods to study sleep. In Fundamentals of sleep and circadian science. Oxford University Press.

Taylor, P. A., Reynolds, R. C., Calhoun, V., Gonzalez-Castillo, J., Handwerker, D. A., Bandettini, P. A., Mejia, A. F., & Chen, G. (2023). Highlight results, don’t hide them: Enhance interpretation, reduce biases and improve reproducibility. Neuroimage, 274, 120138. 10.1016/j.neuroimage.2023.120138

Taylor, P. A., & Saad, Z. S. (2013). FATCAT: (An efficient) Functional and Tractographic Connectivity Analysis Toolbox. Brain Connectivity, 3(5), 523–535. 10.1089/brain.2013.0154

Titone, S., Samogin, J., Peigneux, P., Swinnen, S. P., Mantini, D., & Albouy, G. (2024). Frequency-dependent connectivity in large-scale resting-state brain networks during sleep. European Journal of Neuroscience, 59(4), 686–702. 10.1111/ejn.16080

Tononi, G., & Cirelli, C. (2014). Sleep and the price of plasticity: From synaptic and cellular homeostasis to memory consolidation and integration. Neuron, 81(1), 12–34. 10.1016/j.neuron.2013.12.025

Tu, W., Ma, Z., Ma, Y., Dopfel, D., & Zhang, N. (2021). Suppressing anterior cingulate cortex modulates default mode network and behavior in awake rats. Cerebral Cortex, 31(1), 312–323. 10.1093/cercor/bhaa227

Tu, W., & Zhang, N. (2022). Neural underpinning of a respiration-associated resting-state fMRI network. Elife, 11. 10.7554/eLife.81555

Vernet, M., Quentin, R., Chanes, L., Mitsumasu, A., & Valero-Cabre, A. (2014). Frontal eye field, where art thou? Anatomy, function, and non-invasive manipulation of frontal regions involved in eye movements and associated cognitive operations. Frontiers in Integrative Neuroscience, 8, 66. 10.3389/fnint.2014.00066

Voss, U., & Harsh, J. (1998). Information processing and coping style during the wake/sleep transition. Journal of Sleep Research, 7(4), 225–232. 10.1046/j.1365-2869.1998.00117.x

Watanabe, T., Kan, S., Koike, T., Misaki, M., Konishi, S., Miyauchi, S., Miyahsita, Y., & Masuda, N. (2014). Network-dependent modulation of brain activity during sleep. Neuroimage, 98, 1–10. 10.1016/j.neuroimage.2014.04.079

Wehrle, R., Kaufmann, C., Wetter, T. C., Holsboer, F., Auer, D. P., Pollmacher, T., & Czisch, M. (2007). Functional microstates within human REM sleep: First evidence from fMRI of a thalamocortical network specific for phasic REM periods. European Journal of Neuroscience, 25(3), 863–871. 10.1111/j.1460-9568.2007.05314.x

Wilson, R. S., Mayhew, S. D., Rollings, D. T., Goldstone, A., Przezdzik, I., Arvanitis, T. N., & Bagshaw, A. P. (2015). Influence of epoch length on measurement of dynamic functional connectivity in wakefulness and behavioural validation in sleep. Neuroimage, 112, 169–179. 10.1016/j.neuroimage.2015.02.061

Woo, C. W., Krishnan, A., & Wager, T. D. (2014). Cluster-extent based thresholding in fMRI analyses: Pitfalls and recommendations. Neuroimage, 91, 412–419. 10.1016/j.neuroimage.2013.12.058

Xu, W., De Carvalho, F., Clarke, A. K., & Jackson, A. (2021). Communication from the cerebellum to the neocortex during sleep spindles. Progress in Neurobiology, 199, 101940. 10.1016/j.pneurobio.2020.101940

Yang, F. N., Picchioni, D., de Zwart, J. A., Wang, Y., van Gelderen, P., & Duyn, J. H. (2024). Reproducible, data-driven characterization of sleep based on brain dynamics and transitions from whole-night fMRI. Elife, 13. 10.7554/eLife.98739

Yeo, B. T., Krienen, F. M., Sepulcre, J., Sabuncu, M. R., Lashkari, D., Hollinshead, M., Roffman, J. L., Smoller, J. W., Zollei, L., Polimeni, J. R., Fischl, B., Liu, H., & Buckner, R. L. (2011). The organization of the human cerebral cortex estimated by intrinsic functional connectivity. Journal of Neurophysiology, 106(3), 1125–1165. 10.1152/jn.00338.2011

Zalesky, A., Fornito, A., & Bullmore, E. T. (2010). Network-based statistic: Identifying differences in brain networks. Neuroimage, 53(4), 1197–1207. 10.1016/j.neuroimage.2010.06.041

